# scFTD-seq: freeze-thaw lysis based, portable approach toward high-quality distributed single-cell 3’ mRNA profiling

**DOI:** 10.1101/447524

**Authors:** Burak Dura, Jin-Young Choi, Kerou Zhang, William Damsky, Durga Thakral, Marcus Bosenberg, Joe Craft, Rong Fan

## Abstract

Cellular barcoding of 3’ mRNAs enabled massively parallel profiling of single-cell gene expression and has been implemented in droplet and microwell based platforms. The latter further adds the value for compatibility with low input samples, optical imaging, scalability, and portability. However, cell lysis in microwells remains suboptimal despite the recently developed sophisticated solutions. Here, we present scFTD-seq, a microchip platform for performing <u>s</u>ingle-<u>c</u>ell <u>f</u>reeze-<u>t</u>haw lysis <u>d</u>irectly toward 3’ mRNA <u>seq</u>uencing. It offers format flexibility with a simplified, widely adoptable workflow that reduces the number of preparation steps and hands-on time, with the quality of data and the cost per sample matching that of the state-of-the-art scRNA-seq platforms. Freeze-thaw, known as an unfavorable lysis method resulting in possible RNA fragmentation, turns out to be fully compatible with single-cell 3’ mRNA sequencing, which detects only ~50 bases at the 3’ end. We applied it to the profiling of mixed populations including whole tumors for distinguishing all major cell types and to the profiling of circulating follicular helper T cells implicated in systemic lupus erythematosus pathogenesis. Our results delineate the heterogeneity in the transcriptional programs and effector functions of these rare pathogenic T cells. As scFTD-seq decouples on-chip cell isolation and the following library preparation steps, we envision it to potentially allow the sampling (capture of cells/beads in microwells) at the distributed sites including small clinics or point-of-care settings and downstream processing at a centralized facility, which should enable wide-spread adoption beyond academic laboratories – for any users even with no experience in scRNA-seq library generation.

## INTRODUCTION

Single-cell RNA-sequencing (scRNA-seq) is becoming a mainstay tool in biology research to examine the heterogeneity of complex samples, identify distinct cell subsets, and dissect cell differentiation processes and lineage commitment (1-14). With the recent advances in molecular barcoding techniques and integration of various microfluidic platforms into library preparation steps, it is now possible to sequence thousands of cells with library preparation costs less than $0.1 per cell (15-18).

While many platforms are now available commercially(19-21) or from academia(17,18,22,23) utilizing various approaches, one of the most commonly used technique involves co-isolating each single cell with a uniquely barcoded mRNA capture bead as the enabling step for preparing barcoded libraries. Droplet-based microfluidic techniques have been one of the earliest and widely used approach to achieve such cell-bead co-isolation, encapsulating cells and beads in individual droplets at high-throughputs for massively parallel processing of single-cell transcriptomes (thousands of barcoded cells per run) (17,18,20). However, droplet based techniques has fundamental limitation in cell-bead pairing efficiency, can hardly deal with low input samples (<500), follows an incessant workflow until reverse transcription step, and require major capital or peripheral equipment (for example, 10x Chromium System or home-built DropSeq systems), which limits their portability. As an alternative, microwell arrays have also been adapted for scRNA-seq applications (22-24), and offer several advantages over droplet-based systems including ease of use without bulky equipment, parallelization, compatibility with low-input samples, perturbation assays and imaging cytometry. In addition, microwell arrays also offer format flexibility where they can be used either in closed-environment cell loading format(22) (with a microfluidic channel bonded on top of the array) or in open-surface cell loading format(23) (with no channel on top, loading the array simply using a pipette) depending on the requirements of specific applications.

One of the challenges in using microwell arrays for single-cell isolation and mRNA capture, however, is the cell lysis step where the lysis buffer needs to be introduced into the microwells with negligible material (e.g., mRNA) loss. In recent demonstrations, two approaches are used to address this challenge. In one approach, microwell arrays are used in closed-environment cell loading format, and the reagent loading procedure is automated with a fast flow delivery system (22,24). After cell and bead loading, lysis buffer is introduced followed by immediate sealing of microwells using fluorinated oil. In the other approach, microwell arrays are used in open-surface cell loading format, and the array surfaces are modified prior to use with a specific functionalization chemistry to facilitate reversible sealing with a semipermeable polycarbonate membrane(23). The pores on the membrane enable solution exchange for cell lysis but retain large macromolecules including most mRNAs inside the microwells, thereby preventing mRNA loss. Both approaches have proved successful and paved the way for the use of microwell arrays for scRNA-seq applications. Another one of the challenges is the incessant workflow requiring highly skilled experimentalists to complete multiple lengthy steps in a timely manner (at least until reverse transcription), which is ubiquitous to most scRNA-seq methods. The ability to decouple sample procurement and loading from the sophisticated library generation steps would foster wide adoption by researchers, particularly those in small clinics or in point of care settings who may not have experience in scRNA-seq library generation. Building upon the earlier work and intending to address the remaining issues, in this study we sought to further improve the ease of use of microwell-based scRNA-seq applications by testing the suitability of freeze-thaw cycles as the lysis method. Freeze-thaw, known as an unfavorable lysis method resulting in possible RNA fragmentation, turns out to be fully compatible with single-cell 3’ mRNA sequencing, which detects only ~50 bases at the 3’ end. Compared to detergent or chaotrope based lysis methods, the freeze-thaw method does not initiate lysis immediately as there is no active lysing reagent in the freeze-thaw lysis buffer, and remedies the need for automated rapid fluid exchange or use of a semipermeable membrane. Furthermore, it decouples cell loading and all the following steps, potentially allowing the sampling (capture of cells/beads in microwells) at the distributed sites including small clinics or point-of-care settings and downstream processing at a centralized facility (after shipping) for high quality and consistent preparation of sequencing libraries. As such, scFTD-seq addresses both challenges in a single platform and is potentially enabling for wide-spread adoption of scRNA-seq at further reduced cost through industrialized high volume service for library preparation and sequencing.

In the next sections, we show that our freeze-thaw lysis based approach yields efficient cell lysis and mRNA capture, and high-quality sequencing libraries comparable to those of previously demonstrated massively parallel approaches (17,18,20,22). We named this technique scFTD-seq, as it only requires a <u>f</u>reeze-<u>t</u>haw process to <u>d</u>irectly lyse single cells isolated in bead-containing microwells and capture single-cell-derived mRNAs for transcriptome sequencing. scFTD-seq is compatible with both open-surface and closed-environment cell loading formats, is portable as it does not require any peripheral equipment, and can spare the entire library generation process at the distributed sample procurement sites. We demonstrated the routine practice with <10,000 cells as input, the ability to obtain up to ~5000 single cell transcriptomes per run, less than $150 cost per sample and $0.1 per cell for library preparation, with a performance matching that of the state-of-the-art platforms. We applied it to the profiling of mixed populations including cell lines and whole tumors for distinguishing all major cell types (both tumor and immune cells) and to the profiling of circulating follicular helper T cells and central memory T cells implicated in the pathogenesis of autoimmune disease - systemic lupus erythematosus – in patients. Our results provide insight into the transcriptional and functional heterogeneity of these rare T cells that may underlie their roles in autoimmunity. All these validated scFTD-seq as a high-performance and competent technology for massively parallel single-cell 3’ mRNA sequencing. It may ultimately expand the adoption of scRNA-seq from academic laboratories to any potential users that have no experience in single-cell sequencing library preparation and further harness the power of industrialization to reduce cost and ensure data quality, a key step toward large-scale clinical applications in the future.

## MATERIAL AND METHODS

### Microwell array fabrication and device assembly

Master wafers for microwell arrays were fabricated using SU-8 negative resist. A single layer of resist (SU-8 2035, MicroChem) was spun at 2200-2400 rpm for 30s to yield feature heights of ~50 μm. The wafers were then exposed to ultraviolet light through a transparency mask (CAD/Art Services) to pattern microwells. After developing and baking, wafers were hard baked at 150°C for 30 min, and silanized for 2 h in a vacuum chamber saturated with Trichloromethylsilane (Sigma-Aldrich). Fabrication of microfluidic channels used in closed format operation followed a similar fabrication procedure using SU-8 2075 where channel height was set to ~120 μm (1700-1800 rpm for 30 s).

Devices were made by casting polydimethylsiloxane (PDMS, Sylgard 184, Dow Corning) over the master wafers followed by degassing and curing at 80°C for 6-8 hours. For closed format, both microwell array and microfluidic channel device were set to a final height of 3-4 mm. For open format, microwell array was set to a height of < 0.5 mm by spin coating PDMS to make the array compatible with Agilent clamp (Agilent, G2534A) when bonded to a glass slide. After curing, PDMS was peeled off, and devices were cut to proper sizes to fit on a glass slide. For microfluidic channels, holes for fluidic connections were punctured using a biopsy punch (Miltex, 1.5mm). For open format, microwell arrays were plasma-bonded to a glass slide. For closed format, microwell arrays were first plasma-bonded to microfluidic channel and then to a glass slide.

### scFTD-seq operation – open-surface loading format

For open-surface format, cell and bead loading procedures were adopted largely from SeqWell approach (23) (https://www.nature.com/protocolexchange/protocols/5793#/main). Before cell loading, microwell arrays were plasma-exposed to make the microwell surfaces hydrophilic, and submerged in 1% bovine serum albumin (BSA) in PBS for 30 min for priming. The arrays were then washed with PBS, and solution on top of the array was removed using a pipette just to leave a thin layer of solution. A single cell suspension, 5,000-15,000 cells in 200 μL PBS+1%BSA solution, was then pipetted on the array at multiple positions to cover the entire array area and incubated for 10-15 min with manual rocking of the array intermittently to improve cell loading. After incubation, microwell arrays were washed with PBS to remove unsettled cells. Washing was repeated as many times as necessary to ensure removal of excess cells and confirmed by imaging. Following cell loading, mRNA capture beads (Macosko-2011-10, ChemGenes) were suspended in PBS as 150,000 beads in 100-150 μL, and pipetted onto the array and loaded by gravity into microwells, similar to that described for cells. The array was manually rocked intermittently and we usually obtained loading of the >85-90% of the wells. Excess beads were removed using PBS washes and confirmed by imaging. The freeze-thaw lysis buffer (500 μL in volume; composition: 100 mM Tris-pH 7.5, 10 mM EDTA, 1M NaCl, 5μM DTT, 0.4U/mL Lucigen RNase) was then pipetted on the array and the array was incubated for 5 min at room temperature. The excess freeze thaw lysis buffer was removed leaving ~200 μL of the buffer on top of the array. A glass slide was then used carefully to seal the array (with minimal bead loss, <5%), and secured using a manual clamp (Agilent, G2534A). Three freeze-thaw cycles were performed to lyse the cells, freezing cells in −80°C freezer or dry ice/ethanol bath for 10-15 min and thawing them at room temperature for 10-15 min. Following lysis, microwell array was incubated for an hour inside a wet chamber for mRNA capture onto beads. mRNA binding occurs in the freeze-thaw lysis buffer without the need for buffer exchange. After incubation, the microwell array was unclamped and glass slide was carefully removed. The array was then interfaced with a lifter slip and transferred to a 4 well plate filled with PBS in inverted orientation, similar to described in SeqWell approach(23). The 4-well plated was centrifuged at 1000g for 5 min to release the beads into the PBS solution. Any remaining beads were carefully removed by scraping the array surface gently with a glass slide or a pipette tip. The PBS solution containing the beads were transferred to a 15 mL falcon tube, centrifuged at 1000g for 5 min and transferred to a 1.5 mL Eppendorf tube in 1 mL volume to proceed with reverse transcription.

### scFTD-seq operation – closed-environment loading format

For closed-environment format, microfluidic devices were first infused with PBS and pressurized to remove air bubbles inside the microwells using a manually operated syringe with outlet closed. The devices were filled with 1% BSA in PBS, and incubated at room temperature for 30 min to prevent attachment of cells and molecules on PDMS surfaces. Devices were then washed with PBS prior to cell and bead loading. To establish gravity-driven flow, device outlet was connected to a 10” tubing with a one way stopcock connected at the end while the device inlet is left unconnected to serve as a reservoir. In this configuration, solutions were simply pipetted onto the inlet reservoir and withdrawn into the device through gravity-driven flow by adjusting the height difference between the inlet and the end of the tubing connected to outlet. The one way stopcock further allowed start/stop control over the fluid flow to facilitate cell and bead loading. Similarly, the flow could also be reversed by creating a higher hydrostatic pressure on the outlet side by adjusting the height of the tubing. During scRNA-seq experiments, a single cell suspension, 5,000-10,000 cells in 50 μL PBS+1%BSA solution, was pipetted on the inlet and withdrawn into the device. Once the channel was completely filled with cell solution, the fluid flow was stopped and cells were allowed to settle by gravity. Excess cells were washed out by PBS, and mRNA capture beads, 30,000-120,000 beads in 50-150 μL, were loaded similar to cells. Size exclusion and back-and-forth loading ensured loading of >99% of the microwells with a single bead. Excess beads were washed out with PBS, and 100-200 μL freeze-thaw lysis buffer was introduced into the devices. Fluorinated oil (Fluorinert FC-40), 100-200 μL in volume, was then withdrawn into the devices to seal the microwells. After oil sealing, the tubing at the outlet was disconnected, and the microfluidic devices were exposed to three freeze thaw cycles, 5 min freezing at −80°C freezer or dry ice/ethanol bath and 5 min thawing at room temperature. Following lysis, microfluidic device was incubated for an hour inside a wet chamber for mRNA capture onto beads. mRNA binding occurs in the freeze-thaw lysis buffer without the need for buffer exchange. After incubation, the inlet of the microfluidic device was connected to a syringe filled with 6X saline-sodium citrate (SSC) buffer and the outlet was connected to eppendorf tube with a tubing. The microfluidic device was then inverted and the beads were flushed out of the device into the tube by purging. Centrifugation of the microfluidic device in inverted orientation before purging or gentle tapping on the back of the microfluidic device with a tweezer during purging was used to help move the beads out of the microwells. We were able to recover >95% of the beads using this fashion. Collected beads were centrifuged at 1000g for 1 min, and washed twice with 6X SSC buffer prior to reverse transcription.

### Library preparation and sequencing

Library preparation and sequencing steps follow the same steps as in the DropSeq method (version 3.1 dated 12/28/2015, http://mccarrolllab.com/dropseq/) (17) and SeqWell method(23) where composition of buffers, sequence of primers used, and protocol steps are described in extended detail. Briefly, the beads were first washed with 6X SSC twice after retrieval, and the captured mRNA was reverse-transcribed using Maxima H Minus reverse transcriptase (ThermoFisher) with a custom template switching oligo (AAGCAGTGGTATCAACGCAGAGTGAATrGrGrG). The cDNA coated beads were then treated with Exonuclease I (Exo I, NEB) for 45 min at 37°C to chew away any unbound mRNA capture probes. The beads coated with cDNA was then amplified using a half PCR reaction (i.e., only 13-16 cycles rather than 35-40), using 13 cycles for cell lines or large cells and 16 cycles for primary cells as in SeqWell method(23). The amplified DNA was purified using Ampure XP beads (Beckman Coulter) at 0.6 ratio, and the quality of the amplified DNA was assessed by Agilent BioAnalyzer using high sensitivity chip. Purified cDNA was then pooled and inputted for standard Nextera tagmentation and amplification reactions (Nextera XT, Illumina) using a custom primer instead of i5 index primer to amplify only those fragments that contain the cell barcodes and UMIs. The PCR product was then purified using Ampure XP beads at 0.6X ratio, and the quality of the libraries were checked by Agilent BioAnalyzer high sensitivity chip. The libraries were sequenced on HiSeq 2500 sequencer (Illumina) using a custom primer for Read 1 with 75 cycles on Read 1 and 75 cycles on Read2. For Read1, only the first 20 bases were used in analysis. PhiX libraries were used at 20% as spike-in controls.

### Read alignment and generation of digital gene expression matrix

Transcriptome alignment including barcode/UMI identification and collapsing were performed as described in Dropseq method(23) using DropSeq tools (http://mccarrolllab.com/dropseq). In short, the second read containing the transcript information was trimmed at 5’ end and 3’ end to remove adapter sequence and polyA tail respectively, and labeled with cell barcode and UMI sequences in the first read. The reads were then aligned to reference transcriptome of the corresponding species (mouse, mm10; human, hg19; human-mouse mix, hg19_mm10) using STAR v2.5.2b. The uniquely aligned reads were first grouped by the cell barcode, and then by UMIs with a single-base error tolerance. Total number of distinct UMI sequences for each gene was reported as the number of transcripts corresponding to that gene in the digital gene expression matrix.

### Single-Cell Gene Expression Data Analysis

The data analysis was performed using Seurat(25,26) on the normalized and log-transformed gene expression data. Library size normalization was performed for each cell where transcript numbers for each gene were scaled by the total number of transcripts and multiplied by 10,000.

### Visual confirmation of freeze-thaw lysis

GFP-expressing HUVEC cells (Angio-Proteomie) were grown in EGM-2 MV full medium (Lonza, CC-3202) at 37°C and 5% CO_2_ and trypsinized at >80% confluency. A single cell suspension was prepared and loaded in microwell arrays as described above without the beads. Cells were then exposed to three cycles of freeze-thaw, freezing at −80°C freezer or dry ice/ethanol bath and thawing at room temperature. Representative images of cells before lysis, after one freeze-thaw cycle and three freeze-thaw cycles were taken at randomly chosen locations. Cell lysis was confirmed by distribution of GFP throughout the entire microwell volume. For confirming nuclear lysis, cells were transfected using Cell Light Nucleus-RFP (ThermoFisher) that expresses RFP fused to the SV40 nuclear localization sequence. After incubation overnight, the nuclear staining was confirmed under fluorescence microscope and freeze-thaw experiments were repeated as described above. Nuclear lysis was confirmed by distribution of RFP throughout the entire microwell volume.

### Experiments on freeze-thaw lysis efficiency

Human K562 cells (ATCC) were grown in RPMI medium supplemented with 10% heat-inactivated FBS, 2 mM l-glutamine, 0.1 mM 2-mercaptoethanol, 100 Units/mL penicilin G sodium, and 100 μg/mL streptomycin sulfate. NIH3T3 mouse fibroblasts (ATCC) were cultured in DMEM media containing 10% bovine calf serum, 4 mM L-glutamine and 100 Units/mL penicillin G sodium and 100 μg/mL streptomycin sulfate. For both K562 and NIH3T3, single cell suspensions of 100,000 cells were prepared and lysed in 500 μL volume either using freeze-thaw cycles (in freeze-thaw lysis buffer; 100 mM Tris –pH 7.5, 10 mM EDTA, 1M NaCl, 5μM DTT, 0.4U/mL Lucigen RNase) or detergent-based buffer as used in DropSeq method(17) (100 mM Tris-pH 7.5, 10 mM EDTA, 3% Ficoll PM-400, 0.1% Sarkosyl, 25mM DTT). For freeze-thaw lysis, lysates were prepared for one, two and three freeze-thaw cycles for comparison. Following lysis, mRNA from lysates were isolated using magnetic oligo-dt beads (NEB, S1419S) following manufacturer’s instructions. mRNA was quantified using NanoDrop 2000 spectrophotometer. For qPCR, 10 ng of mRNA was used as input for all conditions and reactions were run on CFX Connect Real-Time PCR machine (Biorad) using SsoFast EvaGreen PCR supermix (Biorad) following manufacturer’s instructions. Primers were obtained from Sigma (KiCqStart primers).

### Species-mixing experiments

Equal numbers of human K562 and mouse NIH3T3 cells were mixed and loaded onto microwell arrays and sequenced at medium depth (average of 20,000-40,000 reads/cell) as described above. Following alignment, cells were filtered based on >10,000 reads, >5000 transcripts and >1000 genes. Cells with >90% transcript alignment to a species-specific transcriptome were identified as belonging to that species. For calculating transcript capture efficiency, cells were sequenced at higher depth (average of 200,000 reads/cell) and reads were downsampled using Picard Tools (broadinstitute.github.io/picard/) to provide appropriate comparisons at different average reads per cell. For comparison to 10X genomics platform, we used the most recent data available at https://support.10xgenomics.com/single-cell-gene-expression/datasets/1.1.0/293t3t3. For comparison to Dropseq, we used the 100 STAMP data (SRA: SRR1748412)) originally published in Macosko et al.(17). For comparison to SeqWell, we used the species mixing 2 data (SRA: SRR5250841) originally published in (23).

For species mixing experiments at 1:10 ratio, K562 cells and NIH3T3 cells were mixed at 1:10 ratio, loaded on the microwell array device and sequenced at shallow sequencing depth (average of 5000 reads/cell) as described. Cells were filtered based on >1000 reads and >500 transcripts. Cells with >90% transcript alignment to a species-specific transcriptome were identified as belonging to that species.

### HEK-K562-HUVEC mixture experiments

HEK, K562 and HUVEC cells were mixed at 1:1:2 ratio, and loaded on the microwell array. Cells were sequenced at shallow depth (average 3500 reads/cell) and filtered based on >1500 reads, >1000 transcripts and >300 genes. Clustering and differential gene expression analysis were performed using the Seurat package(25,26), and cell identities were inferred based on the expression of cell-type specific gene expression.

### Melanoma experiments

For single cell analysis of YUMMER1.7 melanomas, 0.5×10^6 YUMMER1.7-GFP cells were injected subcutaneously into the flanks of 6 week old male C57Bl/6J mice. Tumors were allowed to grow for 21 days; tumor volume was tracked using serial caliper measurement. On day 21, when tumors were roughly 500 mm^3 in volume, mice were euthanized and tumors were harvested. Single cell tumor suspensions were generated by digesting with 1000U/mL collagenase, type IV (Sigma) in RPMI with 2% FBS for 30 mins at 37C with gentle agitation. Cells were then stained using CD45 (eBioscience) and LIVE/DEAD Fixable Far Red Dead Cell stain kit (ThermoFisher). Cells were purified by FACS. Dead cells and debris were excluded and CD45+ cells were enriched in order to increase the frequency of CD45+ cells for sequencing with a final ratio of 1:1 CD45+ and CD45- cells. Cells were sorted into cold RPMI with 10% FBS and kept on ice before processing. For sequencing experiments, cells were loaded onto microwell arrays and sequenced as described above at an average depth of ~25,000 reads per cell. Cells were filtered based on >8000 reads, >1000 transcripts and >300 genes, and analyzed using Seurat package(25,26).

### SLE patient circulating CD4+CXCR5+ T cell experiments

Peripheral blood mononuclear cells (PBMC) were isolated from fresh blood by density-gradient centrifugation on Ficoll-Paque (GE Healthcare), with single cell suspensions stained with the following antibodies: APC-H7-conjugated anti-CD3 (clone SK7), PE-Cy5-conjugated anti-CD45RA (clone HI100), PE-conjugated anti-CD25 (clone M-A251), PE-Cy7-conjugated anti-PD-1 (clone EH12.1), Alexa Fluor 488-conjugated anti-CXCR5 (clone RF8B2) and V450-conjugated anti-CCR7 (clone 150503) (all from BD Bioscience), and Alexa Fluor 700-conjugated anti-CD4 (clone OKT4; from eBioscience). Stained cells were sorted into naive (CD3^+^CD4^+^CD45RA^+^CCR7^hi^ PD-1^lo^;), CXCR5^+^ central memory (Tcm: CD3^+^CD4^+^CD45RA^-^ CXCR5^+^ CD25^lo^ CCR7^high^PD-1^low^), and circulating Tfh (cTfh: CD3^+^CD4^+^CD45RA^-^ CXCR5^hi^ CD25^lo^ CCR7^low^ PD-1^high^) by FACS Aria (BD biosciences), with exclusion of doublets by forward and side scatter.

For sequencing experiments, cells were loaded onto microwell arrays and sequenced as described above at an average depth of ~20,000-40,000 reads per cell. Cells were filtered based on >10000 reads, >1000 transcripts and >400 genes, and analyzed using Seurat package(25,26).

## RESULTS

### scFTD-seq workflow

scFTD-seq utilizes microwell arrays as the platform for co-isolating single cells and uniquely barcoded mRNA capture beads (Figure S1A) prior to freeze-thaw cell lysis. The design of the microwell arrays are similar to the ones described before(22,23), and arrays are made from polydimethylsiloxane (PDMS). The dimensions of microwells are dictated by the size of mRNA capture beads (~35 μm); we used ~45-55 μm in diameter and ~50 μm in height to ensure accommodation of only a single bead as well as most mammalian cell types. This choice of dimensions also facilitates straightforward removal of beads after mRNA capture either by purging (for closed-environment cell seeding) or centrifugation (for open-surface cell seeding) after inverting the devices. The throughput of the microwell arrays is determined by the total number of wells and is highly scalable. In this study, we fabricated arrays with 15,000-70,000 wells to be able to sequence ~1,000-5,000 cells in a single run where the arrays are loaded with a well occupancy rate of 5-10% to minimize dual occupancy (cell duplets). In the open-surface loading format (Figure 1A), the throughput varies but is mainly dictated by the area utilized on the array as well as by the number of cells to be loaded. In the closed-environment loading format (Figure 1A), the throughput can be well controlled by choosing the appropriate channel size and cell loading concentration. We demonstrated up to ~5000 single cells processed per run using a flow channel covering ~30% of a standard 1”x3” glass slide and the scale-up to >15,000 cells per run can be readily achieved without major changes in device design and workflow.

**Figure 1.**
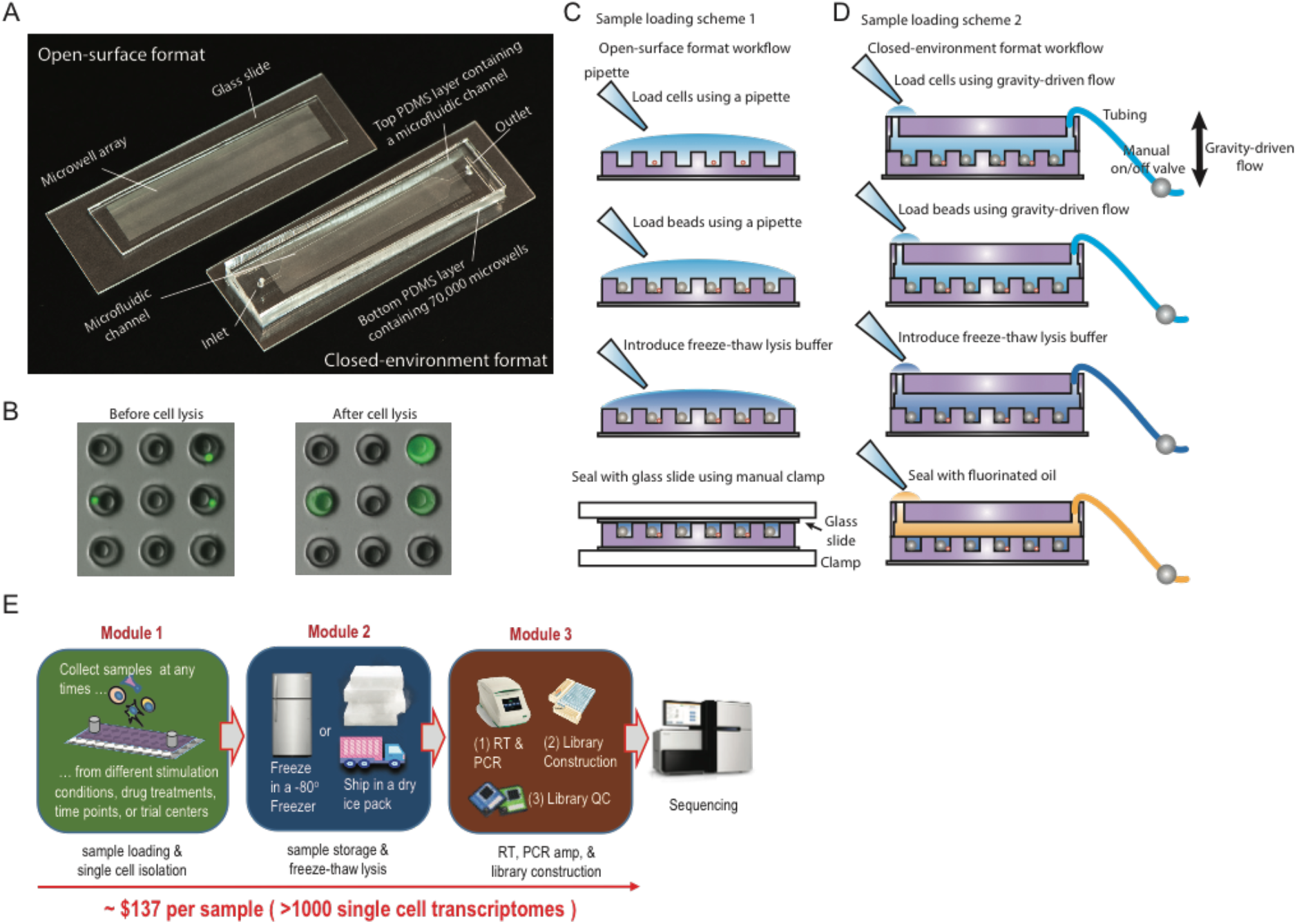
scFTD-seq platform and workflow. (A) Microwell array devices used in scFTD-seq method. For closed-environment format, microwell arrays are bonded to a microfluidic channel. For open-surface format, microwell arrays are used without a channel on top. (B) Cell and bead capture in microwell arrays before (top) and after (bottom) freeze-thaw cell lysis. (C) Open-surface format workflow for scFTD-seq. (D) Closed-environment format workflow for scFTD-seq. (E) Schematic of the scFTD-seq workflow with distributed sampling and modular operation. Library preparation after freeze-thaw lysis follows the protocol described in DropSeq method(17).

To offer format-flexibility for scRNA-seq applications, we established workflows using the microwell arrays both in open-surface and closed-environment loading format (Figure 1C, 1D). In the open-surface format, microwell arrays are used as a miniaturized microtiter plate, and this format offers the advantage of ease of use and potential for co-measurement of mRNA and proteins from the same set of cells, for example, using the well-established single-cell barcode chip(27) or microengraving(28) methods. For scRNA-seq, cells and beads are first loaded on the array sequentially using a pipette and allowed to settle into wells by gravity. This approach usually allows capture of roughly 10-30% of the cells pipetted on the array. Size exclusion ensures each well receives at most a single bead with bead loading efficiencies of >85-90% over the entire array. Following cell and bead capture, a hypertonic solution (herein also called freeze-thaw “lysis” buffer) is pipetted onto the array and the microwells are sealed with a glass slide using a manual clamp. Cell lysis is achieved using three freeze-thaw cycles followed by incubation to allow released mRNAs to be captured onto barcoded beads (Figure 1B). After incubation, glass slide is removed and beads are retrieved by centrifugation (with array in inverted orientation) for subsequent library preparation steps, as described previously(17,23) (Figure 1C, 1E).

In the closed-environment format, the microwell arrays are bonded to a microfluidic channel to introduce cells and reagents using a laminar flow profile (Figure 1A). This format is particularly advantageous due to increased consistency, higher loading efficiency, reduced cell consumption, and minimal waste of barcoded beads. Cell and bead loading time is also shorter compared to open well format as the cells and beads are restrained to a shallow region defined by channel geometry and thickness right on top of microwells, which facilitates the sinking of beads and cells into the wells. Moreover, cells and beads can also be flowed back and forth in the microchannel by reversing the flow to increase capture probability, which makes this format particularly suitable for low-input samples (e.g., <500 cells total) and allows for cell capture efficiencies between ~30-50%. This format is also more conducive for subjecting cells to perturbation assays before transcript capture, which may be useful for many studies such as drug screening applications. With no peripheral fluid handling equipment, we can load cells and beads to this device sequentially using gravity-driven flow, which then settle into wells by gravity (Figure S1B, S1C). Due to the laminar flow profile and the ability to move beads back-and-forth, we routinely achieved >99% bead loading efficiencies with the closed-environment format. The hypertonic buffer is then introduced into the channel followed by sealing with fluorinated oil. As the buffer itself does not initiate lysis, oil sealing can be performed manually using gravity-driven flow with no time constraint or any concern for material loss. Afterwards, the downstream processes including cell lysis, mRNA capture, bead retrieval, reserve transcription, amplification, and library preparation, are performed similar to that described for open-surface format (Figure 1D, 1E).

Importantly, both formats can be operated in a highly-controlled manner with no requirement of any peripheral fluid-handling equipment. Moreover, scFTD-seq workflow is modular, allowing for sample storage after cell/bead loading and pauses at selected steps (Figure S2) without reduction in data quality, which is advantageous for distributed sampling and wide-spread applications. For example, one can use either device to co-isolate cells/beads and freeze down for shipping back to a centralized facility for downstream processing to ensure data quality and consistency (Figure 1E). As such, scFTD-seq combines the most desirable aspects of previously published microwell-based scRNA-seq platforms and offers a more user-friendly workflow with reduced hands-on time, low cost and no need for automated systems or semipermeable membranes (Figure S2, Table S1).

### Technical validation and benchmarking

Repeated cycles of freezing and thawing is an old approach to lyse bacterial cells and has also been demonstrated for mammalian cells including integration in a microfluidic system for gene expression analysis(29). This approach causes formation of ice crystals during freezing and cell expansion upon thawing, eventually leading to rupture of cell membranes. However, it is also known that repeated freeze-thaw can damage large-sized biomolecules and result in fragmentation, an issue for applications such as full-length mRNA sequencing. It was previously utilized to facilitate a detergent-based cell lysis approach in microwells(24), but this method alone has not been considered a viable approach for cell lysis to prepare single-cell mRNA sequencing library. However, we reasoned that the molecular barcoding-based single-cell 3’ mRNA sequencing only detects ~50 bases at the 3’ end, and thus may not be affected by possible mRNA fragmentation induced by freeze-thaw process. Motivated by this reasoning and the previous reports, we explored whether freeze-thaw lysis by itself would lyse the cells efficiently and release their mRNA content sufficiently for single-cell 3’ mRNA transcriptome profiling, and how to optimize this process to achieve the same quality of data.

We first visually confirmed cell lysis by subjecting GFP expressing cells loaded onto microwell arrays to freeze-thaw cycles and imaging the fluorescence of the cell lysate. Even after one freeze-thaw cycle, we observed homogeneous diffusion of cell lysate throughout the wells, which verified the breakage of cell membrane and the ability for large molecules (GFP) to diffuse out from the broken membrane (Figure S3A). To confirm the breakage of the nuclear membrane, we expressed RFP fused to SV40 nuclear localization sequence in cellular nuclei and similarly observed both the cytoplasmic and nuclear lysates homogeneously distribute inside the wells even after a single freeze-thaw cycle (Figure S3B). In addition, we observed little to no change in fluorescence intensity up to 2 h after lysis, indicative of efficient microwell sealing and lysate retention in the microwells (Figure S3C, S3D).

Next, we examined whether freeze-thaw cycles would release sufficient amounts of RNA from cells comparable to detergent based techniques. For this purpose, we compared our freeze-thaw lysis approach to the detergent based approach originally used in DropSeq method(17) in a population based assay. Although we detected higher RNA amounts using detergent based approach compared to freeze-thaw cycles, particularly compared to only one or two cycles, the difference was modest compared with three cycles where the yield was >90-95% of the RNA content as normalized to detergent based approach (Figure S3E, S3G). Critically, when the same amounts of RNA were input for quantitative PCR, we detected similar gene expression levels for a selected set of genes for both human (K562, Figure S3F) and mouse cells (NIH3T3, Figure S3H), confirming the feasibility of freeze-thaw lysis for quantitative gene expression studies with results comparable to those of detergent based techniques.

To validate the overall scFTD-seq workflow and its technical performance, we profiled a mixture of cells from a human leukemic cell line, K562, and a mouse fibroblast cell line, NIH3T3, using our platform in both open-surface and closed-environment loading formats. Bioanalyzer traces of both cDNA and sequencing-ready libraries were very similar to those obtained with detergent based lysis methods (Figure S4). After sequencing, the majority of the cell barcodes uniquely aligned to either the human or mouse transcriptome, suggesting highly species-specific single-cell profiles and minimal cross-contamination between the wells (Figure 2A, 2B, Figure S5). However, open-surface format in general yielded slightly higher number of doublet frequencies (5-7%) compared to closed-environment format, which was due to the less efficient cell loading procedure based on manual pipetting (Figure S6). We detected similar number and distribution of genes and transcripts in both formats in both human (K562) and mouse (3T3) cells, suggesting comparable mRNA capture efficiencies and sequencing performance (Figure 2C).

**Figure 2.**
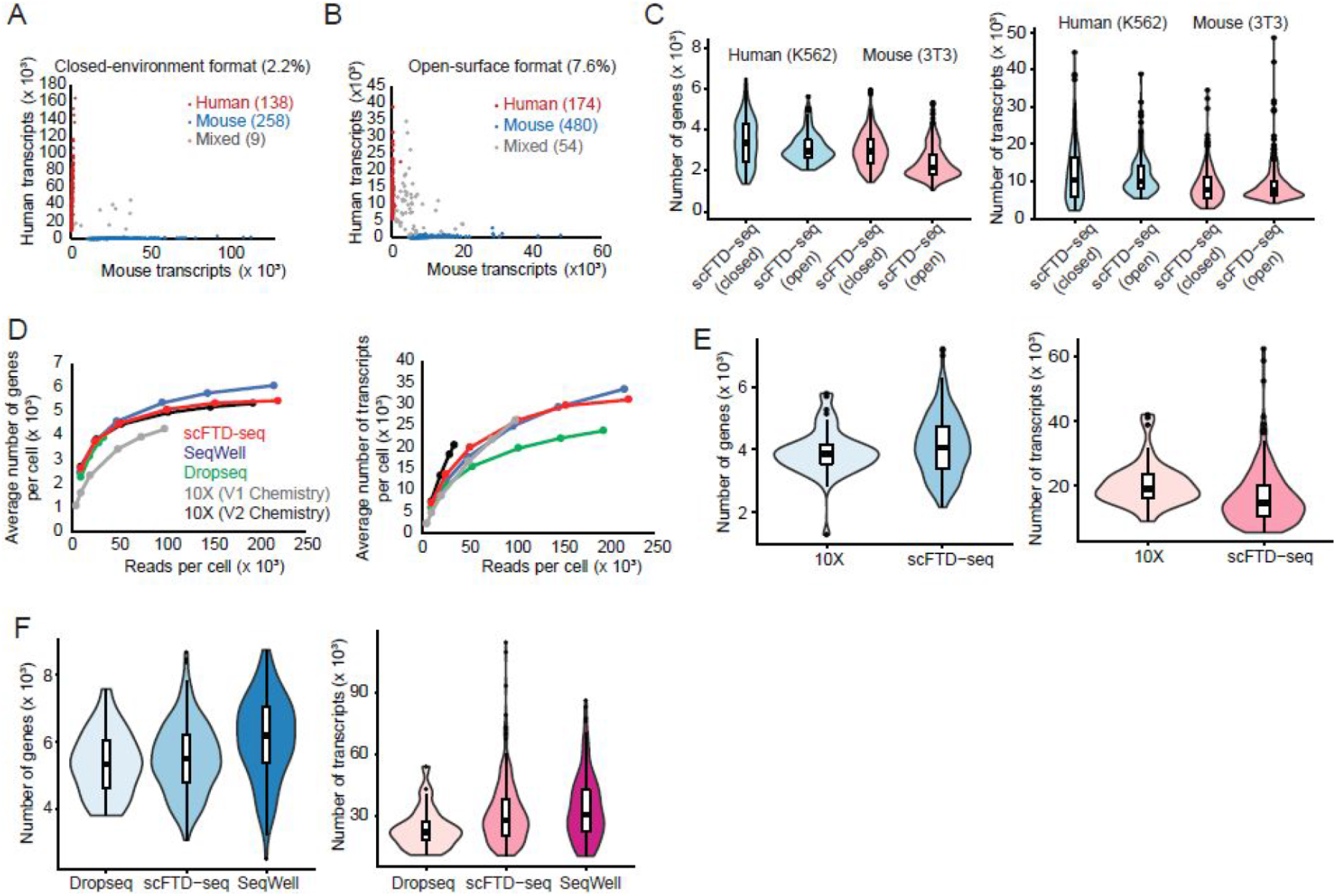
Technical performance of scFTD-seq. (A) Mixed-species experiments reveal single-cell resolution and minimal cross-well contamination for scFTD-seq in closed-environment format (B) Similar results are also obtained for open-surface format but with higher doublet frequency due to the inherent limitation in cell loading. (C) Sequencing performance comparison with 10X Genomics with V1 chemistry (n-538 cells), 10X Genomics with V2 chemistry (n=50 cells), Dropseq (n=27 cells), SeqWell (n=172 cells) and scFTD-seq (n=238 cells) platforms at varying read depths (mouse 3T3 cells). Curves exhibit nearsaturation profiles after 100 million reads per cell. (D) Sequencing performance comparison with 10X Genomics platform (V2). At an average of ~35,000 reads per cell, an average of 5,094 genes and 26,192 transcripts (n=238 cells) are detected in scFTD-seq platform, comparable to 4,233 genes and 27,189 transcripts (n=50 cells) detected in 10X platform in the same cell line (mouse, 3T3). (E) Sequencing performance comparison with Dropseq and SeqWell platforms at saturating read depths using same cell line (mouse, 3T3). An average of 5,469 genes and 31,017 transcripts are detected in scFTD-seq platform (n=238 cells; average of 221,622 reads/cell), comparable to 6,113 genes and 33,586 transcripts (n=172 cells; average of 217,227 reads/cell) in SeqWell, and 5,753 genes and 26,700 transcripts (n=27 cells, average of 194,428 reads/cell) in Dropseq.

We then compared the performance of scFTD-seq with commercially available 10X genomics(20) and state-of-the-art academic platforms(17,23). At varying read depths, we obtained gene and transcript numbers comparable to those of 10X Genomics platform (Figure 2D, 2E). Similarly, we detected an average of ~45,000 transcripts from ~6000 genes in human cells, and ~30,000 transcripts from ~5500 genes in mouse cells at saturating read depths, which was comparable to those of Dropseq and SeqWell platforms (Figure 2D, 2F), including the total number and identity of the genes detected (Figure S7). The sequencing results also revealed high quality libraries where the average fraction of reads mapping to exonic regions were >88% with minimal mapping to ribosomal RNA regions and expected expression percentage of mitochondrial genes (<5%, Figure S8).

To demonstrate the ability for scFTD-seq to process ~5000 single-cell transcriptomes in a single run and to identify cell types in heterogeneous samples, we used a mixture of human (K562) and mouse (NIH3T3) cells mixed at 1:10 ratio, and loaded them onto the largest scFTD-seq platform (70,000 wells). After alignment and filtering, we obtained 5062 single cell transcriptome profiles, 551 K562 cells and 4,511 NIH3T3 cells, which was close to the expected 1:10 mixing ratio (1:8.19, Figure 3A). A similar experiment was also performed using a mixture of three human cell lines, K562, HEK and HUVEC, mixed at 1:1:2 ratio. We recovered 3927 single cell data points after filtering, and unsupervised clustering separated the cells into three major clusters, each of which could be inferred as the corresponding cell type based on the expression of cell-type specific markers (Figure 3B, 3C). We identified 984 K562 cells, 1,000 HEK cells and 1,943 HUVEC cells, which was in good agreement with the expected 1:1:2 mixing ratio.

**Figure 3.**
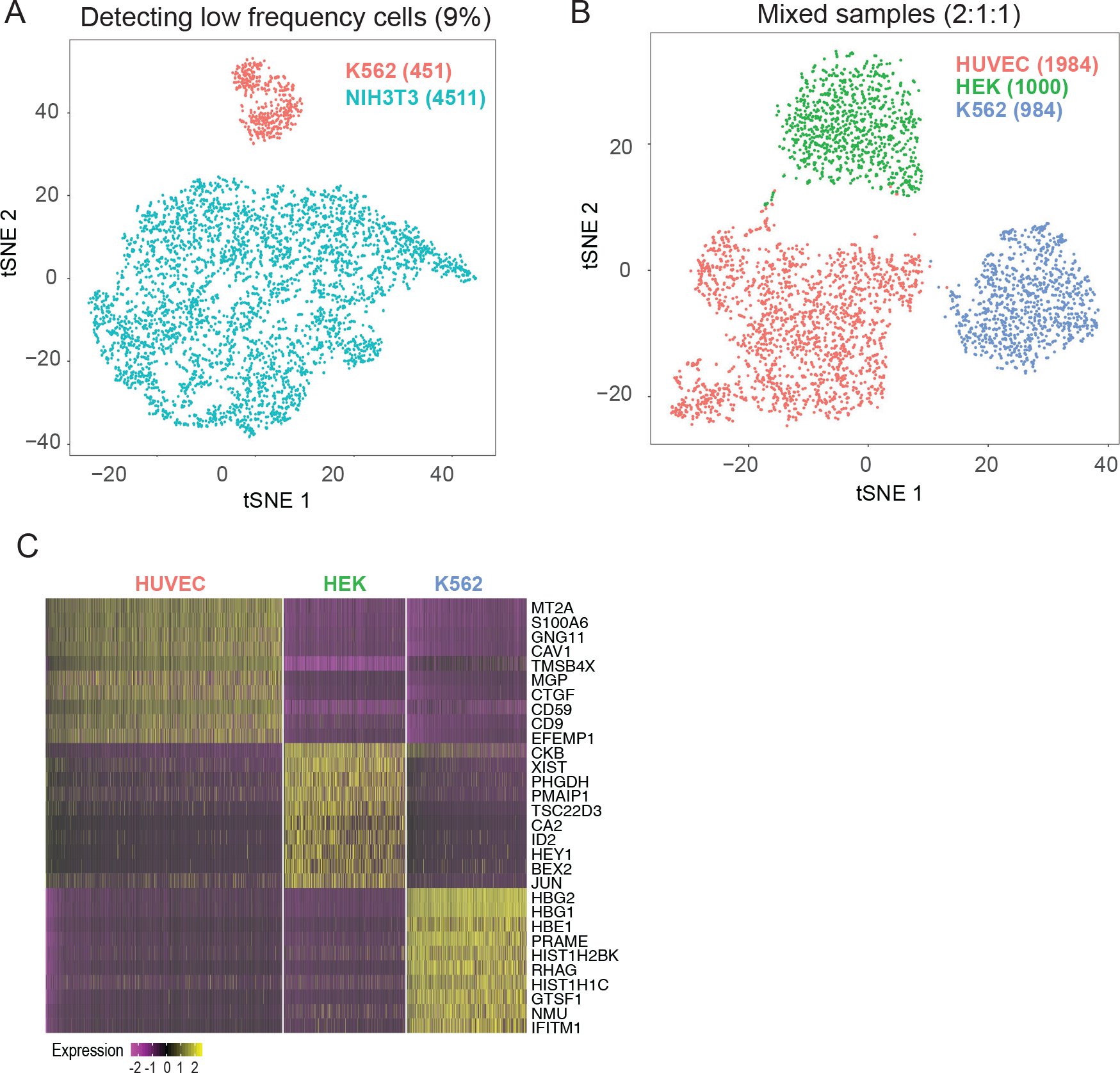
Identifying cell types in mixed samples. (A) t-SNE plot showing the clustering results from a mixture of human K562 and mouse NIH3T3 cells mixed at 1:10 ratio. 5,062 cells were identified after alignment and filtering, of which 451 cells were inferred as human and 4511 as mouse. (B) t-SNE plot showing the sequencing results of a mixture of three human cell lines, HEK-K562- HUVEC, mixed at 1:1:2 ratio. Unsupervised clustering analysis identifies three major clusters. (C) Each cluster is identifiable as the corresponding cell type based on expression of cell-type specific genes.

Collectively, these results demonstrated the feasibility of freeze-thaw cycles as a suitable lysis method to measure single-cell gene expression, and further established scFTD-seq as a robust approach for single-cell 3’ mRNA sequencing with a technical performance matching that of the state-of-the-art platforms.

### Single-cell RNA profiling of whole tumors

To further demonstrate that scFTD-seq can be used with freshly isolated primary cells and enable subpopulation discovery, we profiled cells dissociated from a melanoma tumor generated in the YUMMER1.7 syngeneic mouse model in which tumor cells are also engineered to express GFP for identification(30). After filtering, we obtained 642 cells and detected an average of ~1400 genes and ~4250 transcripts per cell at a depth of ~25k reads/cell. Using clustering analysis, we identified four clusters that formed two distinct major clusters inferred as immune cells (clusters 2, 3, 4) and tumor cells (cluster 1) based on *Ptprc (Cd45)* and *Gfp* gene expression respectively (Figure 4A-4C). Differential gene expression analysis revealed that one of the immune cell clusters (cluster 3) was enriched in the expression of cytotoxicity-associated genes *(Gzmb, Gzmc*, Figure 4D). Examination of canonical lymphocyte markers *(Cd3g, Cd8, Nkg7, Trbc1 and Ctla2a)* further suggested that this cluster was largely composed of cytotoxic T cells and natural killer (NK) cells, whereas the second immune cell cluster (cluster 1) included high numbers of monocytes/macrophages *(Itgam/Cd11b, Cd14, Cd68)* and low numbers of dendritic cells *(Itgax/Cd11c)*, T cells *(Cd3g)* and NK cells *(Nkg7;* Figure S9). The third immune cell cluster (cluster 4) included cells expressing higher levels of complement system genes (C3, C1s, C1ra, Table S2).

**Figure 4.**
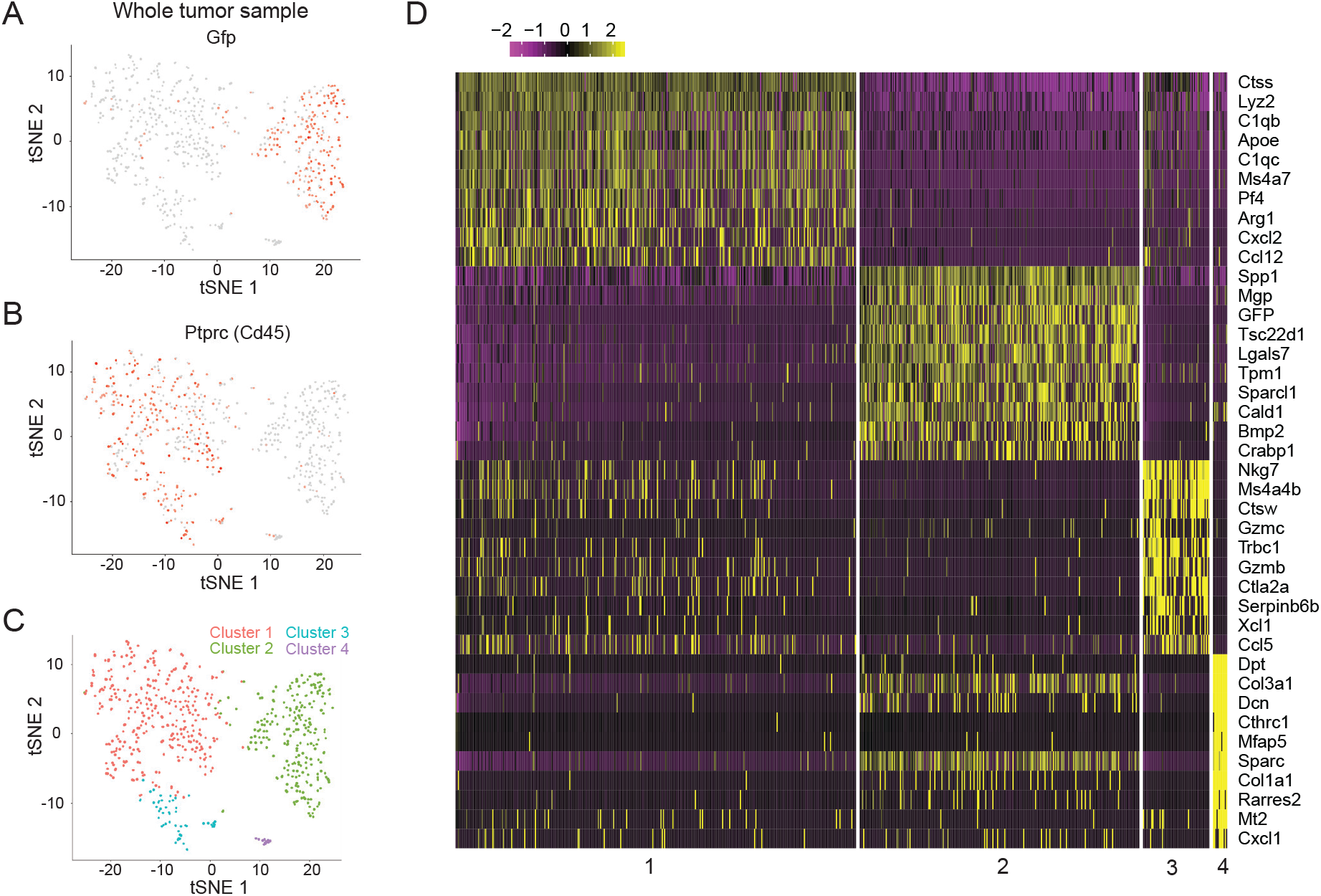
Inferring cell types in heterogeneous samples – whole tumor. (A) t-SNE plot of cells dissociated from melanoma tumor generated in YUMMER1.7 syngeneic mouse model in which tumor cells express *Gfp.* Two major clusters are identified where the left cluster is melanoma cells inferred by *Gfp* expression. (B) Immune cell cluster is inferred by *Ptprc* (*Cd45*) expression. (C) Unsupervised clustering analysis identified 4 clusters where clusters 1 (n=337), 3 (n=56) and 4 (n=12) were identified as immune cells, and cluster 2 (n=237) as tumor cells. (D) Differential gene expression analysis between the identified clusters.

### Single cell RNA profiling of pathogenic T cells from SLE patients

T follicular helper (Tfh) cells are a subset of CD4+ T cells found in secondary lymphoid organs (SLOs) that provide essential help to B cells for their differentiation into plasma and memory cells(31). Evidence from animal models and patients suggests that aberrant functioning of Tfh cells is associated with pathogenic autoantibody production in systemic lupus erythematosus (SLE), an autoimmune disease characterized by immune-complex mediated tissue injury in multiple organs(31). However, investigation of human Tfh cells in SLE has been challenging due to limited access to SLO samples. Studies within the last decade have found that blood CD4^+^CXCR5^+^ cells in humans include a memory compartment of CD4+ T cells that share phenotypic and functional properties with Tfh cells in SLOs, thereby providing a window into the analysis of Tfh cells in SLE(13,32). Further analysis showed that circulating CD4^+^CXCR5^+^ T cells consist of distinct populations with different phenotypes and function, although the combination of markers selected to define these blood Tfh subsets has differed among laboratories(13,32). In our recent work, we used PD-1 and CCR7 markers to divide the CD4^+^CXCR5^+^ population into two subsets, and showed that CD4^+^CXCR5^+^PD-1^high^CCR7^low^ population, termed circulating Tfh-like cells (cTfh), are expanded in SLE patients and have a functional capacity similar to that of Tfh cells in SLO, characterized by higher expression of PD-1 and secretion of IL-21 compared to CD4^+^CXCR5^+^PD1^low^CCR7^high^ central memory T (Tcm) cells(33). Here, we extend our earlier work, applying the scFTD-seq platform to characterize these two cell populations to gain insight into the heterogeneity in their transcriptional programs. In addition, these experiments allowed us to assess the compatibility of scFTD-seq with low-input clinical samples (e.g., a few thousand cells).

After sorting CD4^+^CXCR5^+^PD1^high^CCR7^low^ and CD4^+^CXCR5^+^PD1^low^CCR7^high^ cell populations (from now on referred as cTfh and Tcm cells, respectively, Figure S10A), we divided each sample into two aliquots; the first was activated with ionoymcin and phorbol 12-myristate 13-acetate (PMA) while the second was kept as an untreated control. Ionomycin/PMA stimulation triggers T cell activation pathways, particularly those of cytokine production, and allows probing of a broad array of responses for functional characterization. A small chamber device was used in this study and ~1000 single cells were captured for scFTD-seq. All these samples can be loaded right after sorting but stored for desired time for batch processing to generate sequencing libraries. Here we first analyzed each T cell group individually to seek distinct subpopulations. For untreated samples, although there was considerable cell-to-cell variation, unsupervised clustering did not find any obviously distinguishable clusters (Figure S10B, S10C). For the stimulation condition, clustering identified two major subpopulations for both cTfh and Tcm cells (Figure S10D, S10F). Examining differentially expressed genes, we found that one subpopulation showed stronger activation as inferred from expression of cytokine genes and downregulation of the chemokine receptor genes *CXCR4* and *CCR7*, presumably rendering cells unable to re-enter B cell follicles – an activated phenotype – while the other was not as responsive as inferred from higher *CXCR4* and *CCR7* expression – thus the likely capacity to re-enter follicles – indicating varying levels of responsiveness to even homogeneous stimulation (ionomycin/PMA) within these cell populations (Figure S10E, S10G).

Next, we compared the stimulated and control groups through unsupervised clustering and differential gene expression analysis. For both cTfh and Tcm groups, we identified two major clusters that largely coincided with the stimulated and control groups (Figure 5A, 5D). Differential gene expression analysis showed upregulation of genes related to immune cell signaling and activation - particularly the cytokine genes – along with downregulation of T cell receptor genes (*CD3G, CD247*) and certain chemokine receptor genes (*CXCR4, CCR7*) in the stimulated group (Figure 5B, 5C, 5E, 5F, S11A, S11B), with downregulation of both sets of genes indicating activation. Similarly, gene set enrichment and pathway analyses revealed enrichment in gene sets associated with proliferation, locomotion, cytokine regulation and inflammatory pathways for both stimulated T cell types (Fig. S11D, S11E, Table S3, S4). Notably, while stimulation induced robust cytokine production from majority of the cells, we also observed chemokine gene expression from a small fraction of the untreated cells including *CCL5* and *CCL20* (genes encoding RANTES and MIP-3β, respectively), suggestive of T cells with an activated, inflammatory phenotype circulating in the blood of SLE patients (Figure 5C, 5F). However, upon stimulation, cells exhibited a pronounced polyfunctionality in cytokine gene responses, in that we observed >60% cells with this capacity (Figure S11F-S11I).

**Figure 5.**
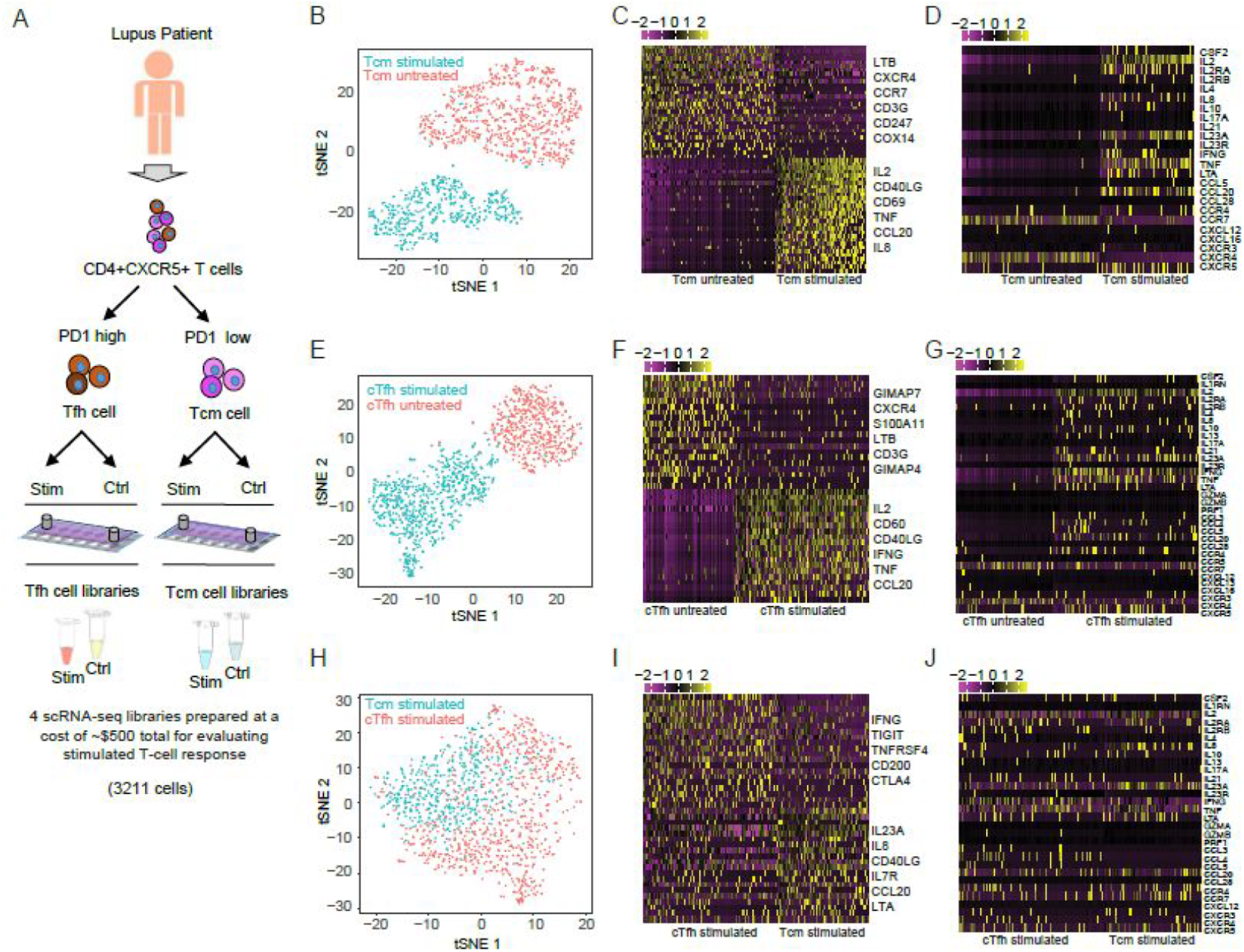
Single cell RNA profiling of circulating CD4^+^CXCR5^+^ T cells and activation states in SLE. (A) Schematic depicting experiment design to profile CD4^+^CXCR5^+^ T cells. (B) t-SNE plot showing the clustering results of stimulated vs untreated CD4^+^CXCR5^+^PD1^low^CCR7^high^ T cells (Tcm). Two major clusters are identified that overlap with stimulated (n=601 cells) and untreated samples (n=946 cells). (C) Differential gene expression analysis of stimulated and untreated Tcm cells. (D) Comparison of cytokine gene expression between stimulated and untreated Tcm cells. (E) t-SNE plot showing the clustering results of stimulated vs untreated CD4^+^CXCR5^+^PD1^high^CCR7^low^ T cells (cTfh). Two major clusters are identified that overlap with stimulated (n=926 cells) and untreated samples (n=738 cells). (F) Differential gene expression analysis of stimulated and untreated cTfh cells. (G) Comparison of cytokine gene expression between stimulated and untreated cTfh cells. (H) t-SNE plot of stimulated Tcm and cTfh cells. (I) Differential gene expression analysis of stimulated Tcm and cTfh cells. (J) Comparison of cytokine gene expression between stimulated Tcm and cTfh cells.

We then analyzed the stimulated cTfh and Tcm groups together to reveal any similarities and differences in their effector functions. We first performed t-distributed stochastic neighbor embedding (t-SNE) analysis where cells with similar transcriptional profiles group together. When we examined the distribution of cTfh and Tcm cells on t-SNE plot, we noticed that the two cell types were separable on the first tSNE component where cTfh cells aggregated largely toward the right while the majority of Tcm cells remained on the left (Figure 5G). This pattern was also observable in principal component analysis, and the two cell types were particularly separable in the first principal component, which consists of selected cytokine and chemokine genes including *IL2, TNF, IFNG* and *CCL20* (*MIP3A*) (Figure S12A, S12B). In agreement with these visual observations, unsupervised clustering separated the cells into three groups in which cluster 1 consisted largely of Tcm cells and clusters 2 and 3 largely of cTfh cells (Figure S12C, S12D). We also captured these differences through differential gene expression analyses and found that, although both activated cell types showed robust upregulation of cytokine genes, their cytokine profiles differed considerably (Figure 5H, 5I, S12E). Particularly, while *IL8, IL23A, TNF* and *CCL20* were largely expressed by the Tcm cells, cTfh cells showed almost exclusive expression of *IL4, IL21, CCL3 (MIPIα), CCL4 (MIP1fβ), CCL5 (RANTES)*, and showed higher expression of *IFNG* and *IL10* (Figure S12F, S13). These results suggest that cTfh cells are a population of heterogeneous effector cells with inflammatory function and are distinct from Tcm cells. We also observed that *IL2* expression followed a bimodal distribution in both cell types with comparable expression levels (Figure S12F). As IL2 transcript and IL-2 protein production by circulating T cells from SLE is decreased compared to healthy subjects, in association with functional changes in T cell receptor mediated activation(34,35), further comparison of these two distinct populations are warranted. Similar to stimulated condition, we also observed differences in baseline gene expression levels between untreated cTfh and Tcm cells (Figure S14).

Overall, these results delineate the transcriptional profiles of cTfh and Tcm cells at the single cell level, and show the intrinsic differences in their effector functions upon stimulation and at the baseline level. A specific expression of CC chemokine family including *CCL3*, *CCL4*, and *CCL5* within the cTfh cells suggests their potential pathogenic role in inflammation via promoting inflammatory responses with possible activation of inflammatory leukocytes.

## DISCUSSION

We have introduced scFTD-seq, a freeze-thaw lysis based single-cell 3’ mRNA-seq approach applicable in microwell array platforms. We validated the technical performance of scFTD-seq with several key demonstrations: (1) using cell lines including the ones used in previous sc-RNAseq publications for head-to-head comparison and mixed samples to access the ability to distinguish heterotypic populations, (2) using primary cells including all types of cells from a whole tumor sample and rare populations of T cells from patients, and (3) examining the activation states of T cells upon stimulation. We have established routine practice with technical performance (throughput, transcript capture efficiency, single-cell resolution, low-input sample compatibility) comparable to state-of-the-art platforms. The cost for barcoding single-cell-derived mRNAs using scFTD-seq (~$150 per sample, mostly the bead cost and Nextera XT kit) is an order of magnitude lower than commercial platforms such as 10x Chromium (>$1500 per sample). Compared to previously established technologies, our platform also offers several key advantages. First, the freeze-thaw lysis eliminates any sophisticated preparatory/follow up steps and any operational challenges, thereby reducing the number of protocol steps and simplifying the overall operation procedure. Second, our platform offers format flexibility, which should enable scRNA-seq applications in a configuration that best suits the demands of the experiment. Importantly, both formats can be operated manually without any additional peripheral equipment, facilitating portability and ease of use. Third, the workflow is modular and can pause at the cell-bead co-isolation step with no effect on the data quality, which reduces the hassle imposed by continuous operation of cell isolation, lysis and reverse transcription, and thereby is more suitable for applications at the distributed sites such as small clinics or point-of-care settings. As such, scFTD-seq presents a reliable and efficient platform with a simplified and modular workflow that can be more readily adopted in both academic, clinical, and large-scale service settings.

We demonstrated the utility of scFTD-seq by applying it to circulating human CD4^+^CXCR5^+^ T cells obtained from SLE patients. Such cells primarily bear a central memory phenotype, with the capacity endowed by CXCR5 and CCR7 expression to migrate to B cell follicles to initiate secondary T-dependent B cell responses(36). Contained within the CXCR5+ pool are also cells with an activated phenotype, with upregulation of PD-1 and downregulation of CCR7, so-called circulating Tfh cells, that may return to the CXCR5-positive memory pool(37). We and others have shown that this subpopulation, identified as CD4^+^CXCR5^hi^PD1^high^ are expanded in the blood of SLE patients(33,37-39). These cells produce IL-21 with their PD-1 expression correlated with disease activity. They expressed lower amounts of CCR7 compared to circulating CXCR5+ central memory cell population, enabling their flow cytometric distinction(33,37).

We have herein extended our earlier work and explored the single cell transcriptional landscape of CD4^+^CXCR5^+^PD1^high^CCR7^low^ T cells (cTfh) in comparison to CD4^+^CXCR5^+^PD1 ^low^CCR7^high^ T cells (Tcm) cells, enabling observations about their heterogeneities and effector function. scRNA-seq identified cells with activation profile (expression of cytokine genes and CD69), even within the untreated control populations despite their low numbers thanks to single-cell resolution. While stimulation strategies are routinely performed to gauge cellular functional capacity, the ability to detect low number of activated cells sensitively by the scFTD-seq technology allows assessing functional states without stimulation, even at the baseline level. Another critical observation was the capacity of the stimulated cell populations, observed in both cell groups, to produce transcripts of multiple proinflammatory cytokines, which if expressed, could contribute to the inflammatory state commonly observed in SLE(40). Such polyfunctionality could also be informative of disease activity, and deserves further investigation. Finally, we observed that, although both cell populations displayed robust cytokine gene expression, their cytokine profiles were distinct with cTfh cells dominantly expressing *IFNG, IL4, IL21, MIPIα, MIP1β and RANTES* - inflammatory cytokine transcripts that could potentially help promote autoreactive B cell activation and induce inflammatory tissue damage. Collectively, these observations provide new insight into functional states of cTfh and Tcm cells that may help explain their role in SLE. Comparison of cellular profiles between patients and healthy subjects is warranted to elucidate mechanisms of SLE pathogenesis and identify potential biomarkers and therapeutic targets.

## ACCESSION NUMBERS

All RNA-seq data will be made available in GEO under accession number GSE109535.

## SUPPLEMENTARY DATA

Supplementary Data are available online.

## ACKNOWLEDGEMENT

Sequencing was performed at the Yale Center for Genome Analysis (YCGA) facility. The molds for microfluidic devices were fabricated in the Yale School of Engineering and Applied Science cleanroom. We graciously thank Michael Power, Chris Tillinghast and James Agresta for their help with fabrication process. We thank Yang Xiao for her help with photography of microfluidic devices and setup.

